# Social rank influences relationships between hormones and oxidative stress in a cichlid fish

**DOI:** 10.1101/2022.10.13.512121

**Authors:** Brett M. Culbert, Shana E. Border, Robert J. Fialkowski, Isobel Bolitho, Peter D. Dijkstra

## Abstract

An individual’s social environment can have widespread effects on their physiology, including effects on oxidative stress and hormone levels. Many studies have posited that variation in oxidative stress experienced by individuals of different social ranks might be due to endocrine differences, however, few studies have evaluated this hypothesis. Here, we assessed whether a suite of markers associated with oxidative stress in different tissues (blood, plasma, liver, or gonads) had social rank-specific relationships with circulating testosterone or cortisol levels in males of a cichlid fish, *Astatotilapia burtoni*. Across all fish, blood DNA damage (a global marker of oxidative stress) and gonadal synthesis of reactive oxygen species (as indicated by NADPH-oxidase (NOX) activity) were lower when testosterone was high. High DNA damage in both the blood and gonads was associated with high cortisol in subordinates, but low cortisol in dominants. Additionally, high cortisol was associated with greater production of reactive oxygen species (greater NOX activity) in both the gonads (dominants only) and liver (dominants and subordinates). In general, high testosterone was associated with lower oxidative stress across both social ranks, whereas high cortisol was associated with lower oxidative stress in dominants and higher oxidative stress in subordinates. Taken together, our results show that differences in the social environment can lead to contrasting relationships between hormones and oxidative stress.

## 1. Introduction

Social interactions have strong effects on the ways that individuals perceive and interact with their environment. While socially dominant individuals use aggression to acquire and maintain priority access to key limited resources—including food, shelter, and access to mates— subordinates may experience social suppression and limited access to these same resources (Clutton-Brock and Huchard, 2013; Milinski and Parker, 1991; Stockley and Bro-Jørgensen, 2011). Consequently, individuals of different social ranks tend to experience different levels of oxidative stress—the relative balance between the production of free radicals (*e.g*., reactive oxygen species (ROS)) and the neutralization of free radicals by antioxidant defenses (Monaghan et al., 2009; Pamplona and Costantini, 2011; Sies et al., 2017)—based on their social rank (Cram et al., 2015; Fialkowski et al., 2022; Losdat et al., 2019; Mendonça et al., 2020; Silva et al., 2018). Dominants tend to experience greater oxidative stress, especially when metabolic demands associated with reproduction and/or territory defense are high (Beaulieu et al., 2014; Cram et al., 2015; Dowling and Simmons, 2009; Georgiev et al., 2015b). However, subordinates may also experience elevated oxidative stress because chronic social suppression can increase ROS production and derail antioxidant function (Beaulieu et al., 2014; Georgiev et al., 2015a; Van de Crommenacker et al., 2011). Since oxidative stress may negatively affect Darwinian fitness (Alonso-Alvarez et al., 2017; Speakman et al., 2015; Stier et al., 2012), understanding how social rank influences oxidative stress may provide more insight into the evolution of social behavior and life history strategies.

Hormones can modulate patterns of oxidative stress by influencing metabolic processes, ROS production, and antioxidant defense/repair mechanisms (Chainy and Sahoo, 2020; Costantini et al., 2011; Darbandi et al., 2018). Levels of androgens (*e.g*., testosterone) and glucocorticoids (*e.g*., cortisol) often vary with social rank (Creel, 2001; Creel et al., 2013; Goymann and Wingfield, 2004; Hirschenhauser and Oliveira, 2006) and both can directly influence levels of oxidative stress (Chainy and Sahoo, 2020; Costantini et al., 2011; Darbandi et al., 2018). While testosterone levels are generally higher in dominants—reflecting higher rates of aggression and/or greater reproductive investment compared to subordinates (Hirschenhauser and Oliveira, 2006; Wingfield, 2017)—rank-based variation in cortisol levels tend to be more variable. For example, glucocorticoid levels can be high in subordinates when they are actively suppressed by dominants, whereas elevated glucocorticoid levels in dominants are thought to reflect the metabolic demands associated with maintaining dominance (Creel, 2001; Creel et al., 2013; Goymann and Wingfield, 2004; Raulo and Dantzer, 2018). Consequently, variation in hormone levels within different social ranks is likely associated with different neural, physiological, and neuroendocrine factors, which could influence relationships between hormone levels and oxidative stress across social ranks. However, most previous studies investigating the relationship between oxidative stress and hormone levels across social ranks have relied on a single measure of oxidative stress and/or within a single tissue (Border et al., 2019; Culbert et al., 2022), which only offers a glimpse into the potential ways that the relationships between hormones and oxidative stress might be modulated by the social environment.

While glucocorticoids tend to increase levels of oxidative stress (Chainy and Sahoo, 2020; Costantini et al., 2011; Spiers et al., 2015), the relationship between androgens and oxidative stress is more complicated (Chainy and Sahoo, 2020; Darbandi et al., 2018). Depending on the tissue, androgens can either increase (Alonso-Alvarez et al., 2007; Chainy et al., 1997; Pansarasa et al., 2002; Zhu et al., 1997) or decrease (Ahlbom et al., 2001; Delgado et al., 2010; Marin et al., 2010; Pang et al., 2002; Pinthus et al., 2007) levels of oxidative stress.

However, most studies which have evaluated relationships between hormones and oxidative stress have used experimental approaches where hormone levels are pharmacologically manipulated (see Chainy and Sahoo 2020; Costantini et al. 2011). While these studies are useful, they can be difficult to interpret biologically because elevated hormone levels in the absence of the activation of upstream regulators of their synthesis (e.g., gonadotropin-releasing hormone for androgens or corticotropin-releasing hormone for glucocorticoids) is unlikely. As well, most studies have used supraphysiological hormone concentrations that may elicit different responses than endogenous levels. Therefore, investigations of the relationship between oxidative stress and endogenous hormone levels, as well as whether this relationship varies with social rank, are needed.

In the present study, we assessed whether the relationship between oxidative stress and hormone levels varied between dominant and subordinate males of the cichlid fish *Astatotilapia burtoni*. These lekking fish live in aggregations where males compete for spawning territories that are used to attract and mate with females (Fernald, 1977). Dominant males that monopolize these territories invest heavily in their reproductive system, having higher levels of androgens (e.g., testosterone) and larger gonads (Fialkowski et al., 2022; Maruska, 2014; Maruska and Fernald, 2011). Conversely, subordinate males that are unable to defend a territory maintain lower testosterone levels, have smaller gonads, and spend most of their time shoaling with females. The costs of being unable to acquire a territory appear high for subordinates because they usually have higher levels of the glucocorticoid hormone cortisol (Border et al. 2019; Carpenter et al. 2014; Fox et al. 1997; Korzan et al. 2014; but see Huffman et al. 2015; Maruska et al. 2013). However, dominant males have greater overall oxidative damage (Border et al., 2019; Fialkowski et al., 2021, 2022)—although this effect is highly tissue-specific—which likely reflects the high metabolic demands associated with reproduction and/or territory defense, as has been suggested in other vertebrates (Beaulieu et al., 2014; Cram et al., 2015; Dowling and Simmons, 2009; Georgiev et al., 2015b). Given these differences in hormone levels and the contrasting physiological demands associated with social rank in *A. burtoni*, we decided to probe whether the relationship between hormone levels and oxidative stress varies across social ranks.

We focused on the blood and plasma because these tissues are often used as a less invasive means of assessing ‘global’ oxidative status in ecological studies (Metcalfe and Alonso-Alvarez, 2010; Monaghan et al., 2009; Speakman et al., 2015). We also evaluated measures of oxidative stress in two tissues that are critical for mediating many of the physiological differences that occur between social ranks. We targeted the gonads because dominant males have larger gonads than subordinate males (Huffman et al., 2012; Maruska and Fernald, 2011) and because they are the primary site for the synthesis of testosterone (Payne and Hales, 2004; Tokarz et al., 2015). As well, we targeted the liver because energetic/metabolic processes in the liver often vary with social rank (Culbert et al., 2019; Gilmour et al., 2012; Kostyniuk et al., 2018) and the liver is one of the primary targets for the metabolic actions of cortisol (Culbert et al., 2021; Faught and Vijayan, 2016; Mommsen et al., 1999).

## 2. Experimental Methods

### 2.1. General Housing

This study took place from June through July of 2017 at Central Michigan University in Mount Pleasant, Michigan, USA. Laboratory-born larvae and juveniles were housed in 110 L tanks until they reached four months old and were moved into 407 L mixed-sex stock tanks that contained gravel substrate and terracotta shelters until experiments started. All tanks were maintained at 28°C with continuous water flow and mechanical/biological filtration. Fish were held under a 12L:12D photoperiod and were fed a mixture of cichlid flakes (Omega Sea LLC) and granular food (Allied Aqua) daily. Prior to the start of experiments, a tagging gun (Avery-Dennison, Pasadena, CA) was used to individually mark focal males with coloured beads affixed to a plastic tag located just below their dorsal fin. All procedures were approved by the Central Michigan University Institutional Animal Care and Use Committee (IACUC protocol 15–22).

### 2.2. Experimental Protocol

All experiments took place in 110 L aquaria that had been separated into 2 compartments using clear, perforated, acrylic barriers. In each compartment (*N* = 31) we placed a group of three males and five females that were between 10 and 12 months old. Each compartment contained one flowerpot shard which was monopolized by the dominant male in each group. In most groups, the largest male maintained social dominance throughout the experiment and hence social rank was experimentally assigned. The initial body mass of fish that became dominant males was 5.2 ± 0.2 g (range of 2.6 6.8 g, *N* = 31) and body mass of focal subordinate males was 4.2 ± 0.1 g (range of 3.0 6.0 g, *N* = 31). Fish could physically interact with members of their own group, as well as interacting with members of the group in the adjacent compartment (see section 2.3. for a description of the behavioural analyses).

Fish were held under these conditions for 6□7 weeks (mean ± SEM: 52.6 ± 0.4 days; range of 48□57 days), after which focal males were terminally sampled. The dominant male and one subordinate male from each tank were rapidly captured and blood (~25□ 100 μL per fish) was collected from the dorsal aorta using a heparinized 26-gauge butterfly needle (Terumo). Blood was transferred to heparinized centrifuge tubes and centrifuged at 4000*g* for 10 minutes. Plasma was separated from the blood cells and both fractions were flash frozen in liquid nitrogen and stored at −80°C. Fish were killed via cervical transection and their mass (to the nearest 0.1 g) and length (to the nearest mm) was recorded. Gonad and liver tissue was collected, flash frozen in liquid nitrogen, and stored at −80°C.

### 2.3. Behavioural Observations

Following a two-week stabilization period, we filmed each group three times a week for four weeks and recorded social behaviours of focal males (the dominant male and one randomly selected subordinate male in each tank) as previously described (Border et al., 2019; Fialkowski et al., 2021). Dominant males were brightly coloured, aggressive, defended a territory and actively courted females, whereas subordinates were visually drab, performed little-to-no aggression, did not court females, and spent large amounts of time shoaling (see Fialkowski et al., 2022). A single observer recorded the number of chases, lateral displays, border displays, courtship displays, and flees that each focal male performed, as well how much time each fish spent shoaling, during a five-minute period. We also calculated a dominance index (# of displays, chases, and reproductive acts □ # of flees) to quantify an individual’s relative dominance. We focused our analyses on the observations taken during the final week because these observations are most relevant to the physiological state of individuals at the time of sampling.

### 2.4. Evaluation of Oxidative Stress

We measured multiple markers of oxidative state that provide different information about antioxidant defenses, ROS production, or overall oxidative stress. To assess antioxidant defenses, we measured total antioxidant capacity (TAC), biological antioxidant potential (BAP), OXY-adsorbent capacity (OXY), and superoxide dismutase activity (SOD). We measured nicotinamide adenine dinucleotide phosphate-oxidase activity (NOX) as an indirect indicator of ROS production. Finally, we assessed oxidative DNA damage by measuring levels of 8-hydroxy-2’-deoxyguanosine (DNA) and used reactive oxygen metabolite levels (ROMs) as a general indicator of oxidative stress. We measured a combination of these measures in the blood (DNA), plasma (ROMs, TAC, OXY, BAP), liver (DNA, NOX, TAC, SOD), and gonads (DNA, NOX, TAC, SOD) of all individuals. For a detailed description of the protocols used to measure these markers, see Fialkowski et al. (2022).

### 2.5. Circulating Hormone Measurement

Plasma cortisol and testosterone levels were measured using enzyme-linked immunosorbent assays (ELISA; Enzo Life Sciences) as previously described (Border et al., 2019). Absorbance was read by a plate reader (Epoch2T, Biotech Instruments, Winooski, VT, USA). Samples were assayed in duplicate (final dilution of 1:30) with mean intra-assay coefficients of variation of 2.9% and 3.3%, and inter-assay coefficients of variation of 2.7% and 2.9% for cortisol and testosterone, respectively.

### 2.6. Statistical Analyses

Statistical analyses were conducted using R (version 4.2.0; R Core Team 2022) and a significance level (α) of 0.05 was used for all tests. Data were visually assessed for normality and homoskedasticity using the ‘performance’ package (Lüdecke et al., 2021). If data did not meet these assumptions, then data were log-transformed prior to analysis. We identified and excluded outliers based on Tukey’s rule (a maximum of two values were excluded per measurement) and calculated either partial eta-squared (□_p_^2^; when multiple dependent variables) or eta-squared values (□^2^; when a single dependent variable) to estimate effect sizes. To investigate whether there were differences in circulating hormone levels (either cortisol or testosterone) between social ranks, we fitted general linear mixed models (LMMs) that included group ID as a random factor using the lmer function in the “lme4” package (Bates et al., 2015). To assess the relationship between levels of testosterone and cortisol we used general linear models (LMs). We then investigated whether measures of oxidative stress were related to circulating hormone levels (either cortisol or testosterone) by fitting LMMs that included hormone levels, social rank (dominant or subordinate), and their interaction as fixed factors and group ID as a random factor. Finally, to test whether the performance of social behaviours was related to circulating hormone levels we used LMs. We analyzed social ranks separately for all analyses that assessed relationships between social behaviours and hormone levels because distributions of social behaviours were highly divergent between dominant and subordinate males. Variation in sample sizes for the different measures of oxidative stress are due to limitations in tissue amount.

## 3. Results

### 3.1. Effects of Social Rank on Circulating Hormone Levels

As previously reported in Fialkowski et al. (2022), circulating testosterone levels of dominants (30.4 ± 1.2 ng mL^-1^) were ~2x those of subordinates (16.8 ± 1.8 ng mL^-1^; χ^2^ = 38.9, □^2^ = 0.47, p < 0.001). In addition, plasma cortisol levels were ~60% higher in subordinates (98.3 ± 13.9 ng mL^-1^) compared to dominants (60.5 ± 12.3 ng mL^-1^; χ2 = 4.08, □^2^ = 0.13, p = 0.04). There was no relationship between cortisol and testosterone levels in either dominants (χ^2^ = 0.13, □^2^ = 0.01, p = 0.72) or subordinates (χ^2^ = 0.16, □^2^ = 0.01, p = 0.69).

### 3.2. Relationships between Oxidative Stress and Testosterone

At the level of the whole animal (*i.e*., blood and plasma), blood DNA damage and plasma testosterone levels showed a negative relationship across both social ranks (p = 0.03; Fig. 1A; Table 1). No other significant relationships between whole animal measures of oxidative stress and circulating testosterone levels were observed (Table 1).

**Figure 1.**
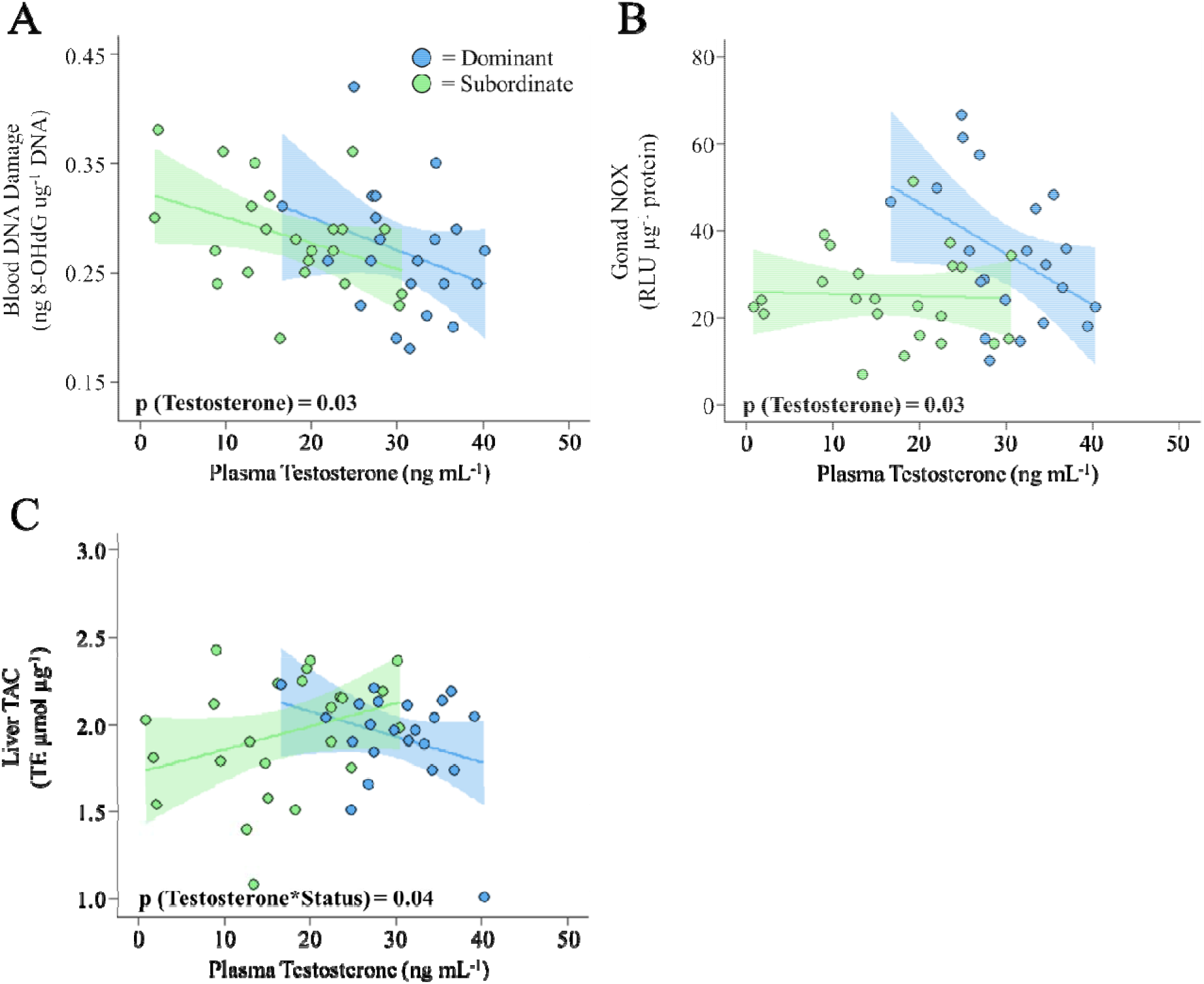
Relationships between plasma testosterone levels and (A) blood DNA damage, (B) gonad NADPH-oxidase activity (NOX), or (C) liver total antioxidant capacity (TAC) in dominant (blue) and subordinate (green) male *Astatotilapia burtoni*. Linear regressions were fit (see text for detailed statistical results) and the shaded area shows the 95% confidence interval of the regression line.

**Table 1.** Relationships between endocrine and oxidative measurements in the blood and plasma of dominant and subordinate *Astatotilapia burtoni* males. Data were analyzed using linear mixed-effects models that included group ID as a random effect. Significant effects (p < 0.05) are indicated in **bold**.

| Tissue | Oxidative Metric | *N*<br>(Testosterone/Cortisol) | Model Term | Plasma Testosterone |  |  | Plasma Cortisol |  |  |
| --- | --- | --- | --- | --- | --- | --- | --- | --- | --- |
| | | | | $\chi^2$ | $\square_p^2$ | *p* | $\chi^2$ | $\square_p^2$ | *p* |
| Blood | DNA Damage<br>(DNA) | Dominants: 21/24 | Hormone | **4.94** | **0.12** | **0.03** | 0.53 | 0.02 | 0.47 |
|  |  | Subordinates: 23/24 | Rank | 0.18 | 0.01 | 0.67 | 2.98 | 0.11 | 0.08 |
|  |  |  | Hormone*Rank | 0.02 | 0.01 | 0.90 | **6.64** | **0.18** | **0.009** |
| Plasma | Reactive Oxygen<br>Molecules (ROMs) | Dominants: 22/24 | Hormone | 0.01 | 0.01 | 0.95 | 2.87 | 0.08 | 0.09 |
|  |  | Subordinates: 24/25 | Rank | 0.50 | 0.01 | 0.48 | 1.59 | 0.05 | 0.21 |
|  |  |  | Hormone*Rank | 0.01 | 0.01 | 0.96 | 0.13 | 0.01 | 0.72 |
|  | Total Antioxidant<br>Capacity (TAC) | Dominants: 21/23 | Hormone | 0.06 | 0.01 | 0.80 | 1.51 | 0.03 | 0.22 |
|  |  | Subordinates: 24 /26 | Rank | 0.71 | 0.02 | 0.40 | 1.74 | 0.04 | 0.19 |
|  |  |  | Hormone*Rank | 1.19 | 0.03 | 0.28 | 0.73 | 0.02 | 0.39 |
|  | Biological Antioxidant<br>Potential (BAP) | Dominants: 13/10 | Hormone | 0.20 | 0.01 | 0.66 | 0.01 | 0.01 | 0.94 |
|  |  | Subordinates: 13 /11 | Rank | 0.02 | 0.01 | 0.89 | 0.43 | 0.03 | 0.51 |
|  |  |  | Hormone*Rank | 0.04 | 0.01 | 0.85 | 0.09 | 0.01 | 0.76 |
|  | OXY-Absorbent<br>Test (OXY) | Dominants: 11/17 | Hormone | 0.60 | 0.03 | 0.44 | 0.01 | 0.01 | 0.96 |
|  |  | Subordinates: 14 /17 | Rank | 0.14 | 0.01 | 0.71 | 0.57 | 0.02 | 0.45 |
|  |  |  | Hormone*Rank | 0.45 | 0.02 | 0.50 | 0.05 | 0.01 | 0.83 |

In the gonads, NOX activity was negatively associated with circulating testosterone levels across all fish (p = 0.03; Table 2); however, this pattern was more evident in dominants (Fig. 1B). For liver TAC values, a significant interaction between social rank and plasma testosterone concentrations was detected (p = 0.04; Table 2), indicating that a rank-specific relationship was present. In dominant males, TAC values were higher when testosterone was low, whereas TAC values were higher when testosterone was high in subordinate males (Fig 1C). No other significant relationships between circulating testosterone levels and measures of oxidative stress in the gonads or liver were observed (Table 2).

**Table 2.**
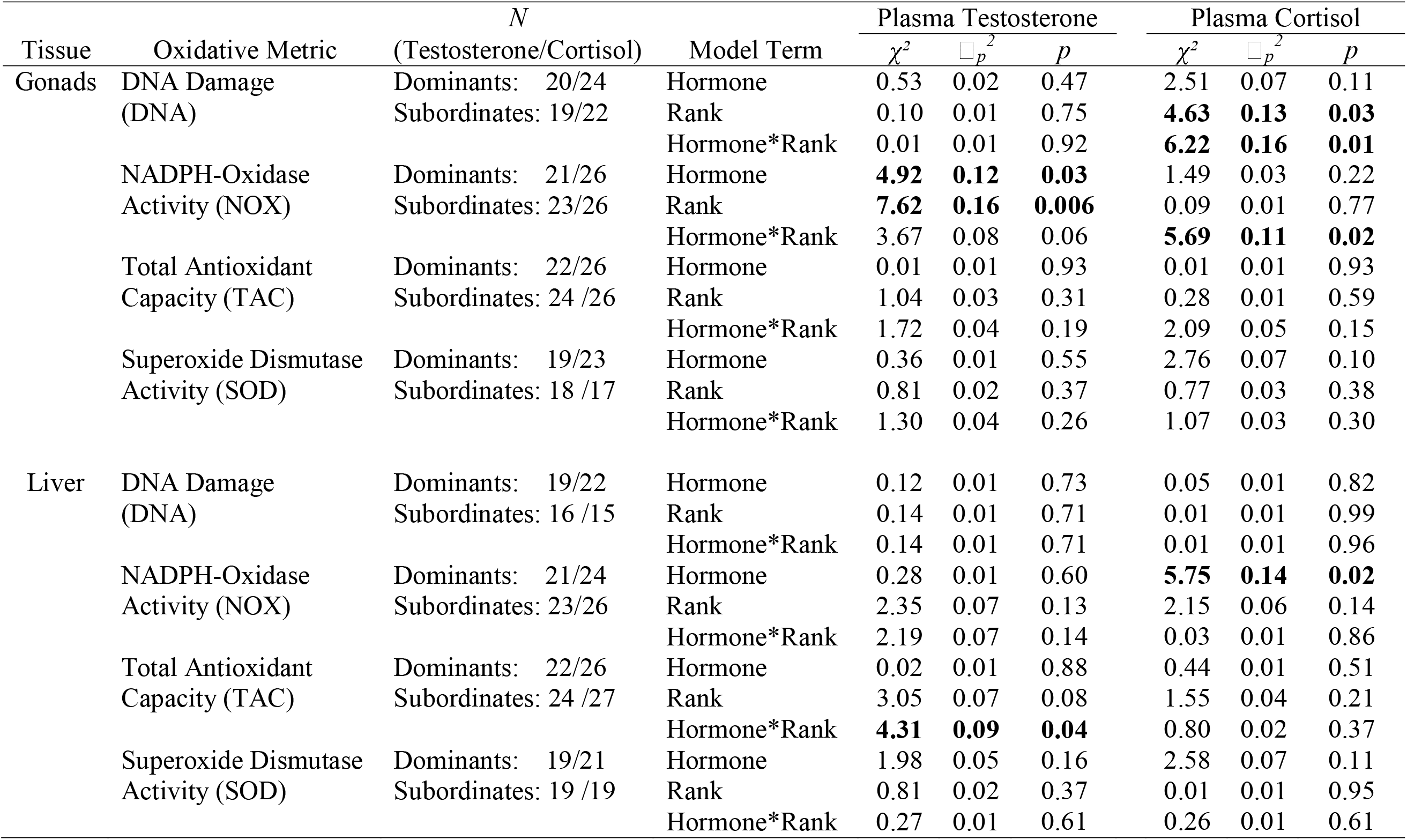
Relationships between endocrine and oxidative measurements in the gonads and liver of dominant and subordinate *Astatotilapia burtoni* males. Data were analyzed using linear mixed-effects models that included group ID as a random effect. Significant effects (p < 0.05) are indicated in **bold**.

### 3.3. Relationships between Oxidative Stress and Cortisol

At the level of the whole animal, blood DNA damage was significantly affected by the interaction between social rank and plasma cortisol concentrations (p = 0.009; Table 1). Blood DNA damage was higher when cortisol levels were low in dominant males, while blood DNA damage was higher when cortisol levels were high in subordinate males (Fig. 2A). No other significant relationships between whole animal measures of oxidative stress and circulating testosterone levels were observed (Table 1).

**Figure 2.**
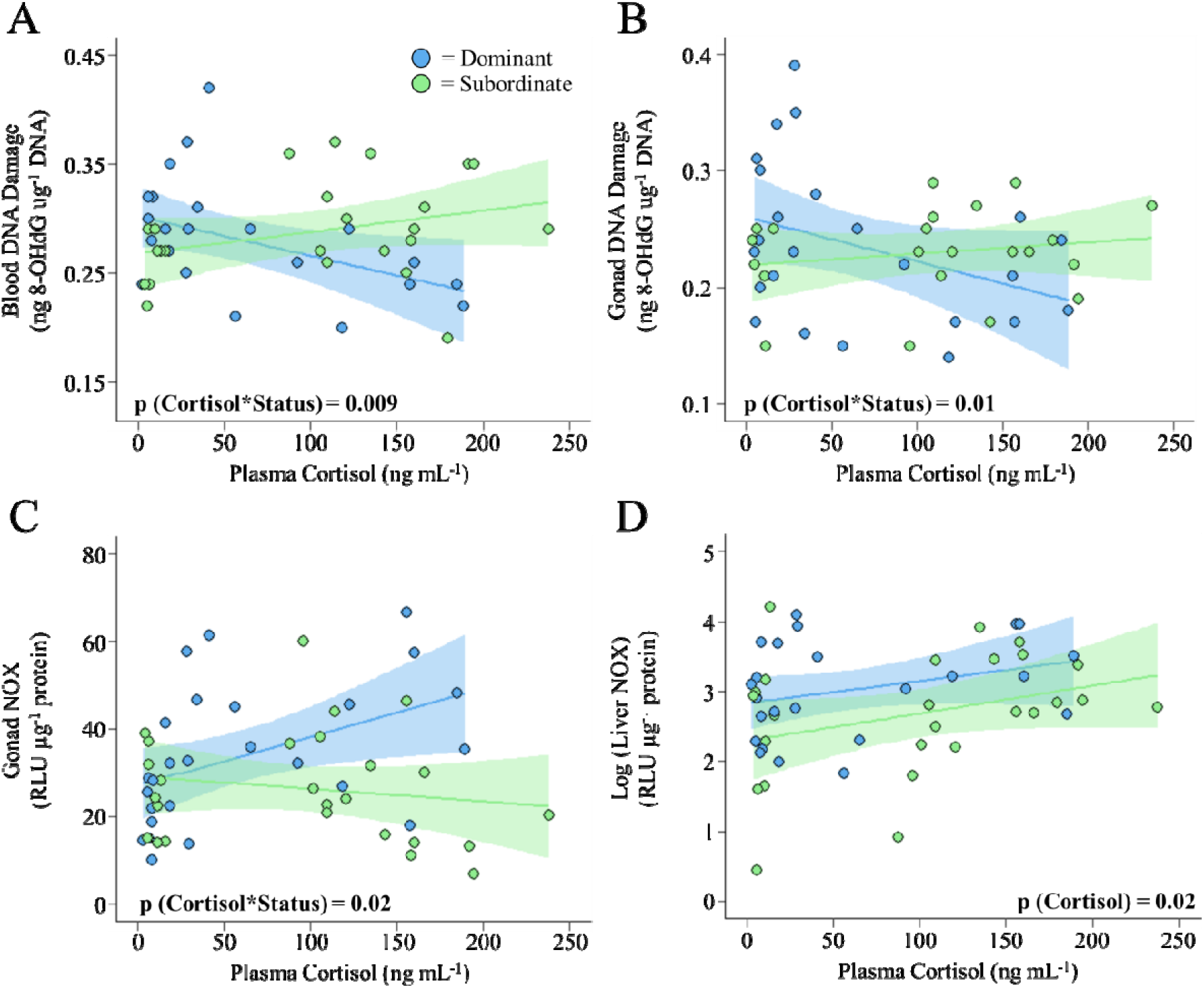
Relationships between plasma cortisol levels and (A) blood DNA damage, (B) gonad DNA damage, (C) gonad NADPH-oxidase activity (NOX), or (D) liver NADPH-oxidase activity (NOX) in dominant (blue) and subordinate (green) male *Astatotilapia burtoni*. Linear regressions were fit (see text for detailed statistical results) and the shaded area shows the 95% confidence interval of the regression line.

Gonad DNA damage was also significantly affected by the interaction between social rank and plasma cortisol concentrations (p = 0.01; Table 2), whereby DNA damage was higher when cortisol was low in dominants versus when cortisol was high in subordinates (Fig. 2B). In the gonads and liver, NOX activity was also related to cortisol levels (Table 2). In the gonads, NOX activity was significantly affected by the interaction between social rank and plasma cortisol concentrations (p = 0.02; Table 2) with NOX activity increasing with cortisol levels in dominant males and exhibiting a slight decrease as cortisol levels increased in subordinate males (Fig. 2C). Liver NOX activity increased with cortisol concentrations in individuals of both social ranks (p = 0.02; Table 2; Fig. 2D). No other significant relationships between circulating cortisol concentrations and measures of oxidative stress in the gonads or liver were observed (Table 2).

### 3.4. Relationships between Social Behaviours and Circulating Hormone Levels

The number of times that dominant males attempted to court females had a significant positive relationship with circulating levels of both testosterone (F = 4.46, □^2^ = 0.18, p = 0.048) and cortisol (F = 6.38, □^2^ = 0.21, p = 0.02). No other significant relationships between hormone levels and social behaviours were detected for either social rank (Supp. Tables 1 & 2).

## 4. Discussion

In the current study, we tested whether *in vivo* relationships between oxidative stress and circulating levels of testosterone or cortisol varied between dominant and subordinate cichlids. We found that high levels of oxidative stress (high DNA damage in blood and a high capacity for ROS synthesis (NOX activity) in the gonads) tended to be associated with low testosterone levels, irrespective of social rank. In contrast, high levels of oxidative stress (DNA damage in the blood and gonads) were associated with high cortisol levels in subordinate males and low cortisol levels in dominant males. Additionally, the capacity for ROS synthesis (NOX activity) in the liver and gonads of dominant males was greater when cortisol levels were high, suggesting that dominant males with high cortisol could prevent DNA damage despite high rates of ROS production in at least some tissues. Overall, our data suggest that relationships between oxidative stress and hormones are affected by an individual’s social rank, as well as the tissue in which oxidative measures are assessed.

The relationship between androgens and oxidative stress is complex. While androgens can stimulate ROS production—likely reflecting, at least in the part, the stimulatory effect that androgens have on metabolic rate (Buchanan et al., 2001; Fernando et al., 2010; Ros et al., 2004)—they can also activate protective antioxidant mechanisms (Chainy and Sahoo, 2020; Darbandi et al., 2018). Consequently, exogenous treatment with androgens can either reduce oxidative stress (Ahlbom et al., 2001; Delgado et al., 2010; Marin et al., 2010; Pang et al., 2002; Pinthus et al., 2007) or increase oxidative stress (Alonso-Alvarez et al., 2007; Chainy et al., 1997; Pansarasa et al., 2002; Zhu et al., 1997) depending on the tissue studied and/or concentration used. However, fewer studies have evaluated relationships between oxidative stress and endogenous androgen levels. Recently, Mentesana and Adreani (2021) reported no relationship between plasma testosterone levels and circulating concentrations of OXY, ROMs, or glutathione peroxidase in wild rufous horneros (*Furnarius rufus*). While we also found no relationship between plasma testosterone and circulating OXY or ROMs, we observed lower levels of global DNA damage (*i.e*., in the blood) in individuals with high testosterone levels, irrespective of their social rank. Therefore, in addition to highlighting the importance of evaluating multiple measures of oxidative stress in several tissues, our data appear to be consistent with previous reports that testosterone reduces oxidative damage—potentially by increasing antioxidant defenses (see Delgado et al., 2010). However, the only evidence of a positive relationship between antioxidant defenses and testosterone was observed in the liver of subordinates, and the opposite pattern occurred in dominant males. Thus, the negative relationship between testosterone and global DNA damage does not appear to be driven by widespread upregulation of antioxidant defenses. Instead, testosterone may be helping to limit DNA damage by reducing gonadal NOX activity (especially in dominants), and although it is unclear the extent to which gonadal ROS production might influence global DNA damage, the observed negative relationship between testosterone and NOX activity has previously been observed (Juliet et al., 2004; Tam et al., 2003). Alternatively, given that high levels of oxidative stress can have inhibitory effects on gonadal androgen synthesis (Glade and Smith, 2015; Kopalli et al., 2022; Turner and Lysiak, 2008), high NOX activity in the gonads (and the associated increase in ROS production) may have supressed testosterone synthesis. This would help to explain why the negative relationship between gonadal NOX activity and testosterone levels was stronger in dominant males since the scope for reducing testosterone synthesis would be greater in dominant males.

Hepatic and gonadal NOX activities (an indicator of ROS production) had a positive relationship with cortisol levels, especially in dominant males. These results are consistent with previous reports that high glucocorticoid levels cause oxidative stress (Costantini et al., 2011; Spiers et al., 2015), likely by stimulating ROS production via direct actions on NOX activity (Espinoza et al., 2017; Kračun et al., 2020; Seo et al., 2012; You et al., 2009) or by causing general mitochondrial dysfunction (Mitsui et al., 2002; Oshima et al., 2004). Because NOX generates large amounts of ROS, high levels of glucocorticoids (Flaherty et al., 2017) and/or elevated NOX activity is usually associated with higher rates of DNA damage (Polytarchou et al., 2020; Weyemi and Dupuy, 2012). However, while a positive relationship between cortisol and DNA damage was observed in both the blood and gonads of subordinate males, DNA damage in the blood and gonads of dominant males was lower when cortisol levels were high. It is likely that these social rank-specific patterns reflect the different ecological/social factors associated with elevated cortisol levels in dominant males compared to subordinate males. While elevated cortisol levels in subordinate males typically reflects social suppression, high cortisol levels in dominant males are thought to reflect the steep metabolic demands associated with maintaining dominance (Creel et al., 2013; Dantzer and Newman, 2022). Here, we detected a positive relationship between cortisol levels and how many times dominant males attempted to court females, suggesting that high cortisol in dominant male *A. burtoni* may primarily be driven by the metabolic demands associated with reproduction. Proximately, the divergent relationships between cortisol levels and DNA damage in dominant males versus subordinate males could relate to dominant male *A. burtoni* having greater levels of ROMs in their plasma compared to subordinate males (Border et al., 2019; Fialkowski et al., 2021, 2022). Chronic exposure to ROMs can upregulate mechanisms that protect and repair DNA (Santa-Gonzalez et al., 2016; Van Houten et al., 2018), and future studies should assess whether such rank-related differences occur in *A. burtoni*. Alternatively, this difference might reflect rank-specific differences in the relative susceptibility of DNA to oxidative damage and/or the efficiency of DNA repair mechanisms, potentially owing to variation in epigenetic markings (e.g., methylation state; Chen and Zhu 2016; Gong and Miller 2019) between dominant and subordinate males (Hilliard et al., 2019; Snyder-Mackler et al., 2019; Tung et al., 2012).

Dominant males that were actively courting females had higher levels of both testosterone and cortisol. This is unsurprising because testosterone is often associated with reproduction and mating (Hirschenhauser and Oliveira, 2006; O’Connell and Hofmann, 2012; Pankhurst, 2016), and—as previously discussed—the high metabolic demands associated with reproduction are thought to be a major contributor to glucocorticoid levels in dominant individuals (Creel et al., 2013; Dantzer and Newman, 2022). While many studies have evaluated the relationship between oxidative stress and reproduction (Alonso-Alvarez et al., 2017; Blount et al., 2016; Costantini, 2018; Speakman et al., 2015; Stier et al., 2012), there have been only a handful of investigations into the potential oxidative costs associated with courtship and no consistent relationship has emerged (see Liao et al., 2018; Montoya et al., 2016; Simmons et al., 2018). In the current study, we did not detect any relationships between measures of oxidative stress and courting behaviour in dominant male *A. burtoni* (data not shown). These results suggest that, while both courtship and oxidative stress are potentially influenced by cortisol and testosterone levels, the proximate mechanisms underlying these relationships likely differ given the lack of direct relationship between courtship and oxidative stress. However, additional work is required to determine the proximate basis for these differences.

In conclusion, our study provides one of the first comprehensive evaluations of how the social environment influences relationships between hormones and measures of oxidative stress across several tissues. Since oxidative stress (Costantini, 2018; Monaghan et al., 2009; Speakman et al., 2015) and hormone levels (Alonso-Alvarez et al., 2009; Crespi et al., 2013) can both influence life-history strategies, it is important to understand how oxidative stress, hormone levels, and the social environment interact with one another. Additional work is required to determine the specific factors responsible for the rank-specific relationships observed between markers of oxidative stress and hormone levels in the current study; especially for cortisol since a greater number of rank-specific relationships with oxidative stress were detected for cortisol versus testosterone. For instance, manipulations of resource availability and/or altering levels of intraspecific competition would help to determine which ecological factors might be driving the contrasting relationships between dominants and subordinates. Overall, this work represents an important step in understanding the role(s) that ecological and social factors have on the relationship between hormones and oxidative stress.

## Data accessibility

Supporting data can be found in the attached supplemental file.

## Competing Interests

The authors declare no competing interests.

## Funding

This work was supported by a Faculty Research and Creative Endeavors grant of Central Michigan University to PDD, and a CMU graduate research grant to SEB. BMC was supported by a Doctoral Canadian Graduate Scholarship (CGS-D) provided by the Natural Sciences and Engineering Research Council of Canada (NSERC) and an Ontario Graduate Scholarship (OGS).

## Acknowledgments

We would like to thank Hannah Janeski, Deric Learman, and Benjamin Swarts for technical assistance.

---

**Supplemental Table 1.** Relationships between plasma testosterone levels and social behaviours in dominant or subordinate *Astatotilapia burtoni* males. Data were analyzed using linear models and significant effects (*p* < 0.05) are indicated in **bold**. Note that displays and courts could not be analyzed for subordinate males and flees could not be analyzed for dominant males because none were performed.

| Behaviour | Dominant Males |  |  | Subordinate Males |  |  |
| --- | --- | --- | --- | --- | --- | --- |
| | *F* | $\eta^2$ | *p* | *F* | $\eta^2$ | *p* |
| Displays Performed | 0.54 | 0.03 | 0.47 | NA | NA | NA |
| Chases Performed | 0.51 | 0.02 | 0.48 | 0.73 | 0.03 | 0.40 |
| **Courts Performed** | **4.46** | **0.18** | **0.048** | NA | NA | NA |
| Flees Performed | NA | NA | NA | 0.09 | 0.01 | 0.77 |
| Dominance Index | 2.11 | 0.10 | 0.16 | 0.01 | 0.01 | 0.94 |
| Time Spent in Shoal (s) | 2.04 | 0.09 | 0.17 | 1.06 | 0.05 | 0.31 |

**Supplemental Table 2.** Relationships between plasma cortisol levels and social behaviours in dominant or subordinate *Astatotilapia burtoni* males. Data were analyzed using linear models and significant effects (*p* < 0.05) are indicated in **bold**. Note that displays and courts could not be analyzed for subordinate males and flees could not be analyzed for dominant males because none were performed.

| Behaviour | Dominant Males |  |  | Subordinate Males |  |  |
| --- | --- | --- | --- | --- | --- | --- |
| | *F* | $\eta^2$ | *p* | *F* | $\eta^2$ | *p* |
| Displays Performed | 2.51 | 0.09 | 0.13 | NA | NA | NA |
| Chases Performed | 0.10 | 0.01 | 0.76 | 2.01 | 0.07 | 0.17 |
| **Courts Performed** | **6.38** | **0.21** | **0.02** | NA | NA | NA |
| Flees Performed | NA | NA | NA | 0.03 | 0.01 | 0.85 |
| Dominance Index | 0.23 | 0.01 | 0.63 | 0.18 | 0.01 | 0.68 |
| Time Spent in Shoal (s) | 2.36 | 0.09 | 0.14 | 0.51 | 0.02 | 0.48 |

## Notes

### Competing Interest Statement

The authors have declared no competing interest.

